# Reporter gene assay for membrane fusion of extracellular vesicles

**DOI:** 10.1101/2021.02.16.431359

**Authors:** Masaharu Somiya, Shun’ichi Kuroda

## Abstract

**Abstract:** Extracellular vesicles (EVs) secreted by living cells are expected to deliver biological cargo molecules, including RNA and proteins, to the cytoplasm of recipient cells. There is an increasing need to understand the mechanism of intercellular cargo delivery by EVs. However, the lack of a feasible bioassay has hampered our understanding of the biological processes of EV uptake, membrane fusion, and cargo delivery to recipient cells. Here, we describe a reporter gene assay that can measure the membrane fusion efficiency of EVs during cargo delivery to recipient cells. When EVs containing tetracycline transactivator (tTA)-fused tetraspanins are internalized by recipient cells and fuse with cell membranes, the tTA domain is exposed to the cytoplasm and cleaved by protease to induce tetracycline responsive element (TRE)-mediated reporter gene expression in recipient cells. This assay (designated as **E**V-mediated **t**etraspanin-**t**TA **d**elivery assay, ETTD assay), enabled us to assess the cytoplasmic cargo delivery efficiency of EVs in recipient cells. With the help of a vesicular stomatitis virus-derived membrane fusion protein, the ETTD assay could detect significant enhancement of cargo delivery efficiency of EVs. Furthermore, the ETTD assay could evaluate the effect of potential cargo delivery enhancers/inhibitors. Thus, the ETTD assay may contribute to a better understanding of the underlying mechanism of the cytoplasmic cargo delivery by EVs.

## Introduction

Extracellular vesicles (EVs) are secreted by living cells and contain biomolecules derived from the donor cells. The physiological role of EVs remains largely unknown and they were formerly known as the “garbage bin” of cells for excretion of the unwanted molecules or organelles. Several studies have shown the cellular disposal role of EVs ^1,2^ although a vast majority of current EV research focuses on the cargo delivery of EVs. Since EVs contain cargo proteins and RNAs, their contents can be transferred from a donor cell to a recipient cell via a paracrine or endocrine mechanism. Recently, EV-mediated cargo delivery events in pathophysiological settings, such as cancers, have attracted considerable attention. Several studies have reported that EVs are involved in tumor suppression ^3,4^ and tumor progression ^5,6^. Several studies have demonstrated that EVs can deliver small RNAs to recipient cells and elicit phenotypic changes. However, there is limited evidence that demonstrates cargo delivery by EVs into recipient cells ^7^. Many confounding factors in the experimental conditions and contaminants in the EV fraction ^8^ must be taken into account in the cargo delivery experiments, to draw a conclusion on “EV cargo transfer hypothesis” ^9^.

The main challenge in current EV research is the lack of a feasible and reliable assay to evaluate the functional cargo delivery process in the recipient cells ^9,10^. Several reporter assays that demonstrate the functional delivery of cargo proteins or RNAs have been reported, including miRNA ^11,12^, Cre-LoxP ^13,14^, and CRISPR/Cas9-gRNA reporters ^15^. However, these assays are influenced by various confounding factors including non-EV components in the EV fraction. Although the readout of these assays is informative for deciphering the delivery mechanism of EVs in recipient cells, a more precise reporter assay is needed. Mechanistically, cytoplasmic cargo delivery should occur after endocytosis and subsequent membrane fusion, or direct fusion with the plasma membrane ^16^. Upon membrane fusion, the luminal side of EVs is exposed to the cytoplasm of recipient cells and release their cargo. The functional delivery assay should reflect the biological delivery mechanism, especially membrane fusion of EVs.

In this study, we developed a reporter assay to quantify the membrane fusion of EVs in recipient cells. In this assay, following fusion of EVs with the cell membrane of the recipient cells, a transcription factor is released from the EVs and then upregulates the expression of a reporter gene (luciferase or fluorescence protein). This assay provides a biologically orthogonal readout and enables us to accurately interpret the cargo delivery process of EVs.

## Materials and Methods

### Materials

The chemical reagents and antibodies used in this study are listed in Supplementary Table 1. All NanoLuc substrates were purchased from Promega. The plasmids used in this study are listed in Supplementary Table 2 and deposited at Addgene. Plasmids were constructed using PCR-based methods (Gibson Assembly ^17^) and confirmed by Sanger sequencing.

### Cell culture and transfection

Human embryonic kidney HEK293T cells (RIKEN Cell Bank) were maintained in 10% (v/v) fetal bovine serum (FBS)-containing Dulbecco’s modified Eagle’s medium (DMEM) supplemented with 10 μg/mL penicillin-streptomycin solution. Cells were cultured at 37°C under 5% CO_2_ in humidified conditions.

Transfection of HEK293T cells was performed as follows: cells were plated in a cell culture dish or multi-well plate and cultured overnight. The next day, the cells were transfected using 25-kDa branched polyethyleneimine (PEI, Sigma). The ratio of plasmid DNA to PEI was 1: 4 (weight). After 24–96 h, the cells were used in the subsequent experiments. Cell culture supernatant was collected after 2–4 days and centrifuged at 1,500 ×g for 5 min to remove cell debris.

### NanoLuc assay

To quantify the expression level of the reporter NanoLuc, the transfected cells were lysed and mixed with NanoLuc substrate (Nano-Glo Luciferase Assay System; Promega) according to the manufacturer’s instructions. Luminescence signal from the cell lysate was measured by using a plate reader, Synergy 2 (BioTek).

### Characterization of tTA-fused proteins in cell lysate and EVs

Protein expression was assessed by western blotting. Briefly, lysates of the transfected cells (total protein was extracted using radioimmunoprecipitation assay [RIPA] buffer [Nacalai Tesque] containing a protease inhibitor cocktail [Nacalai Tesque]) or the supernatant was mixed with reductant-free sample buffer and incubated at room temperature for 20 min. Proteins were separated by sodium dodecyl sulfate polyacrylamide gel electrophoresis (SDS-PAGE) and transferred to polyvinylidene difluoride (PVDF) membrane. Proteins on the membrane were detected using antibodies (Supplementary Table 1) and ImmunoStar LD reagent (FUJIFILM Wako Pure Chemical). As a loading control for cell lysates, the membrane was probed with anti-GAPDH antibody.

### Concentration of EVs

EVs were concentrated by PEG precipitation. The supernatant was mixed with 4× PEG solution (40% PEG 6000 [w/v], 1.2 M NaCl, 1 × PBS [pH 7.4]), and kept at 4°C overnight. The next day, the supernatant was centrifuged at 1,600 ×g for 60 min to pellet the EVs. After decantation, the pellet was resuspended in PBS. Typically, 5–10 mL of the supernatant was concentrated to 100–200 μL.

### Reporter assay

For the membrane fusion reporter assay, recipient HEK293T cells (10^4^ cells/well in 96-well plate) were transfected with plasmids encoding tobacco etch virus (TEV) protease (TEVp) and TRE3G-NlucP (PEST motif-fused NanoLuc [NlucP]^18^ under tetracycline responsive element [TRE] promoter), and cultured overnight. The next day, the recipient cells were treated with donor culture supernatant or concentrated EVs and further incubated at 37°C for up to 26 h. To assess the effect of various compounds on membrane fusion efficiency, recipient cells were treated with the compound 1 h before the addition of supernatant or EVs. After incubation (2–26 h), the expression of NanoLuc in the recipient cells was measured as described above.

Reporter expression in recipient cells was also evaluated using an enhanced green fluorescent protein (EGFP) gene. Recipient cells (10^4^ cells/well in 96-well plate) transfected with pTetOn-EGFP (EGFP under TRE promoter) and pcDNA3.1-TEVp were treated with EVs, and then observed under a fluorescence microscope IX70 (Olympus) after 24 h. Cre recombinase-based reporter assay was performed in the same way; recipient HEK293T cells were transfected with reporter plasmid (encoding LoxP-flanked mKate and EGFP under the CMV promoter) and plasmid encoding TEVp, treated with EVs for 24 h the following day, and then observed under a fluorescence microscope.

### Statistical analysis

Data were analyzed using Student’s *t*-test or one-way ANOVA following either *post hoc* Tukey’s HSD or Dunnett’s tests. Statistical analysis was performed using the Real Statistics Resource Pack software created by Charles Zaiontz.

## Results

### Characterization of tTA-fused tetraspanins

To establish a reporter assay that can measure the membrane fusion of EVs, we first prepared plasmids encoding human tetraspanins CD9, CD63, or CD81 with C-terminal fusion of the TEVp cleavage site, followed by tTA (Fig. 1A). As shown in Fig. 1B, tTA-fused tetraspanin is cleaved in the presence of TEVp and releases the transcription activator tTA. When the EVs containing tTA-fused tetraspanin are internalized and fused with the endosomal membrane, luminal tTA are exposed to the cytoplasmic side, and TEVp in the recipient cells cleaves the TEVp site, followed by cytoplasmic release of tTA and induction of the reporter gene expression under the TRE promoter (Fig. 1C). We designated this assay the **E**V-mediated **t**etraspanin-**t**TA **d**elivery (ETTD) assay.

**Fig. 1.**
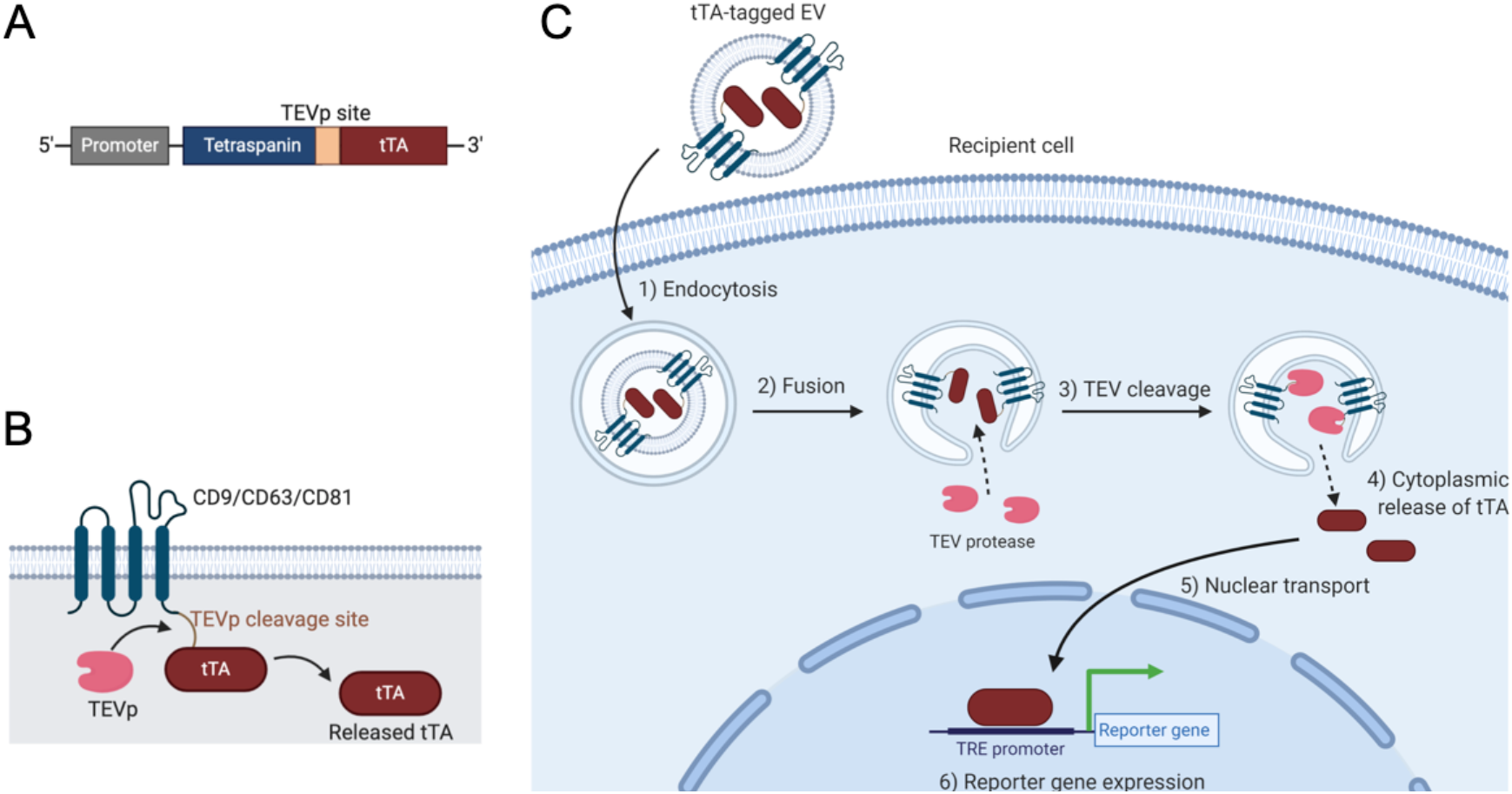
Summary of the ETTD assay. (A) Schematic representation of tTA-fused tetraspanin. Tetraspanin and tTA flank a TEVp recognition site. (B) Topology of tTA-fused tetraspanin protein. Upon the cleavage by TEVp, tTA is released from membrane-anchored tetraspanin. (C) Schematic representation of the ETTD assay. EV containing tTA-fused tetraspanin is taken up by cells by endocytosis (1), and fuses with the endosomal membrane (2). After cleavage by cytoplasmic TEVp (3), tTAs are released into the cytoplasm (4). Released tTAs are transported to nucleus (5), and induce expression of reporter gene under TRE promoter (6).

To demonstrate the feasibility of the above system, HEK293T cells were transfected with plasmids encoding tTA-fused tetraspanin with or without plasmid encoding TEVp. As shown in Fig. 2A, the expression of tTA-tetraspanins in the cell lysate was confirmed by western blotting. In the presence of TEVp, tTA was cleaved and released from the tTA-fused protein. While CD9 and CD81 showed obvious tTA bands, CD63 showed only a weak band in the absence of TEVp and no band in the presence of TEVp. This is probably due to low expression of CD63 in HEK293T cells compared to CD9 and CD81. When HEK293T cells were transfected with both NlucP (under the TRE promoter) and tTA-fused proteins, co-expression of TEVp strongly induced Nluc expression (Fig. 2B), suggesting that tetraspanin-anchored tTA was unable to translocate into the nucleus, and therefore could not induce reporter gene expression. In contrast, expression of non-fused tTA protein continually induced reporter gene expression regardless of the co-expression of TEVp. These results suggest that tTA-fused tetraspanins induce reporter gene expression in the recipient cells only when the cells express TEVp.

**Fig. 2.**
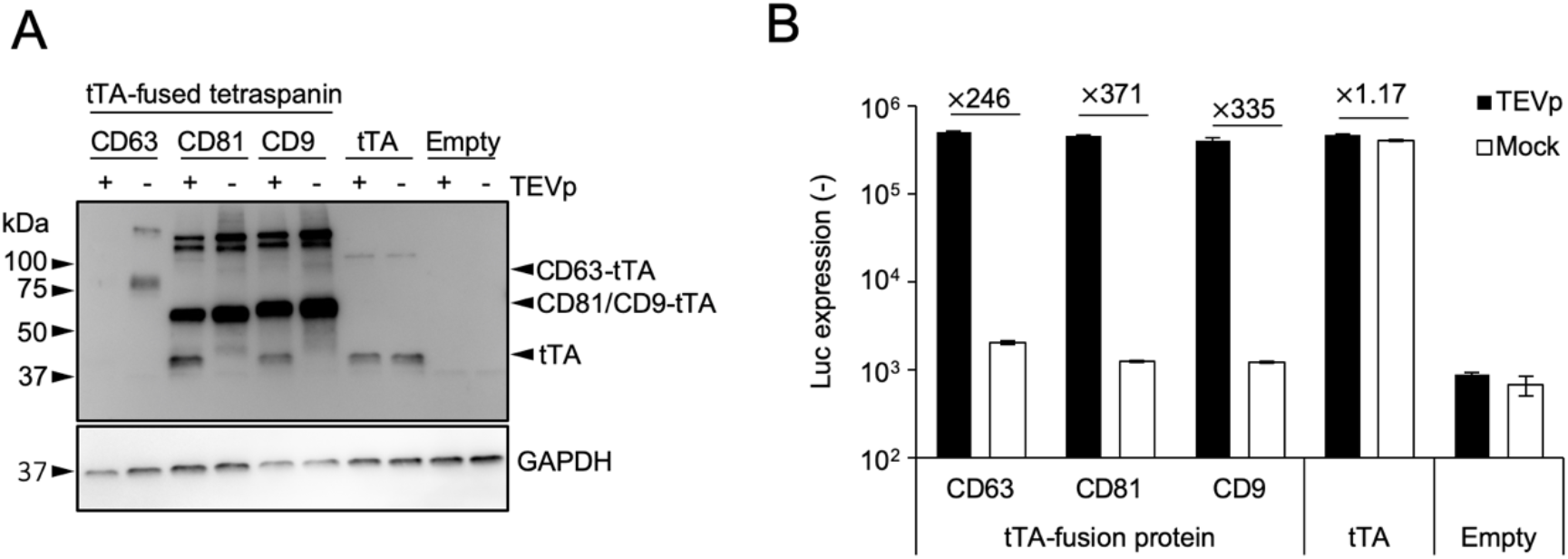
Characterization of tTA-fused tetraspanins. (A) Cells were transfected with plasmids encoding tTA-fused tetraspanins and TEVp. After 48 h, cells were lysed and subjected to western blotting. Upper and lower panels represent immunoblotting using anti-TetR antibody and anti-GAPDH antibody, respectively. The expected molecular weights based on the amino acid sequences were as follows: CD63-tTA, 63.2 kDa; CD81-tTA, 63.4 kDa; CD9-tTA, 63.0 kDa; tTA, 36.9 kDa. (B) Expression of NanoLuc under TRE3G promoter in HEK293T cells co-expressing tTA-fused tetraspanins and TEVp. As controls, plasmids encoding tTA without tetraspanin fusion and empty expression plasmid were used. Numbers above the bars indicate the fold increase in NanoLuc expression compared to the mock transfection. N = 3, mean ± SD

To characterize the tTA-fused tetraspanins (CD81 and CD9) in EVs, supernatants from transfected HEK293T cells were concentrated by PEG precipitation and analyzed by western blotting (Fig. 3A and 3B). Both tTA-fused CD81 and CD9 were detected with anti-CD81 and CD9 antibodies, respectively. The tTA-fused proteins were also detected with an anti-TetR antibody, indicating that the released EVs contain full-length tTA-fused CD81 or CD9. As a control for the ETTD assay, vesicular stomatitis virus glycoprotein (VSV-G) was co-expressed in donor cells, as VSV-G is known to strongly facilitate membrane fusion and subsequent cargo delivery of EVs ^19–21^. VSV-G was detected in the EV fraction, strongly suggesting that released EVs display VSV-G on their surface along with tTA-fused tetraspanins.

**Fig. 3.**
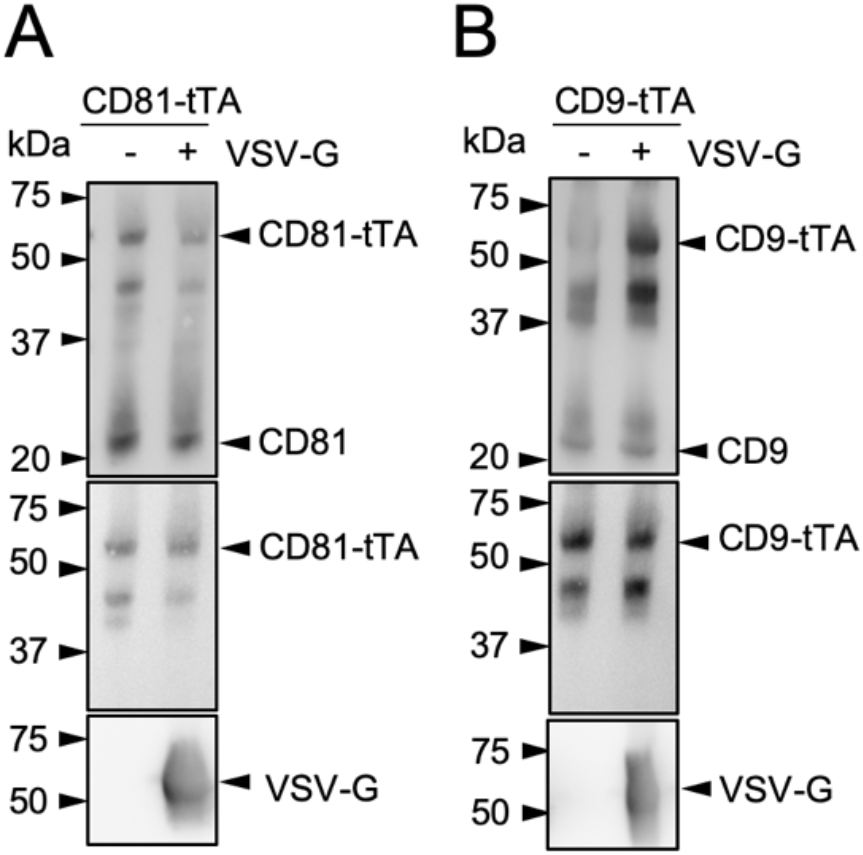
Characterization of HEK293T-derived EVs containing tTA-fused tetraspanins by western blotting. (A) EVs containing CD81-tTA with or without VSV-G. Antibodies used were as follows; top, anti-CD81 antibody; middle, anti-TetR antibody; bottom, anti-VSV-G antibody. (B) EVs containing CD9-tTA with or without VSV-G. Antibodies used were as follows; top, anti-CD9 antibody; middle, anti-TetR antibody; bottom, anti-VSV-G antibody. The expected molecular weights based on the amino acid sequences were as follows: CD81-tTA, 63.4 kDa; CD9-tTA, 63.0 kDa; VSV-G, 57.7 kDa.

### Validation of ETTD assay for cargo delivery of EVs

We first attempted to assess whether the unconcentrated cell culture supernatant from donor cells was capable of inducing reporter gene expression in recipient cells. As shown in Fig. 4A, treatment of recipient cells with donor supernatant containing tetraspanin-tTA fusion protein induced reporter gene expression only when the donor cells were transfected with virus-derived fusogenic protein VSV-G. This result suggested that the concentration process is not necessary to evaluate EV membrane fusion in the ETTD assay if the EVs possessed potent fusogenic activity. While the supernatant containing tTA-fused CD81 and CD9 induced > 10-fold increase in NanoLuc expression, the supernatant containing tTA-fused CD63 showed less induction (up to 5-fold). This may reflect the lower expression level of tTA-fused CD63 in the donor cells compared to CD9 and CD81 (Fig. 2A).

**Fig. 4.**
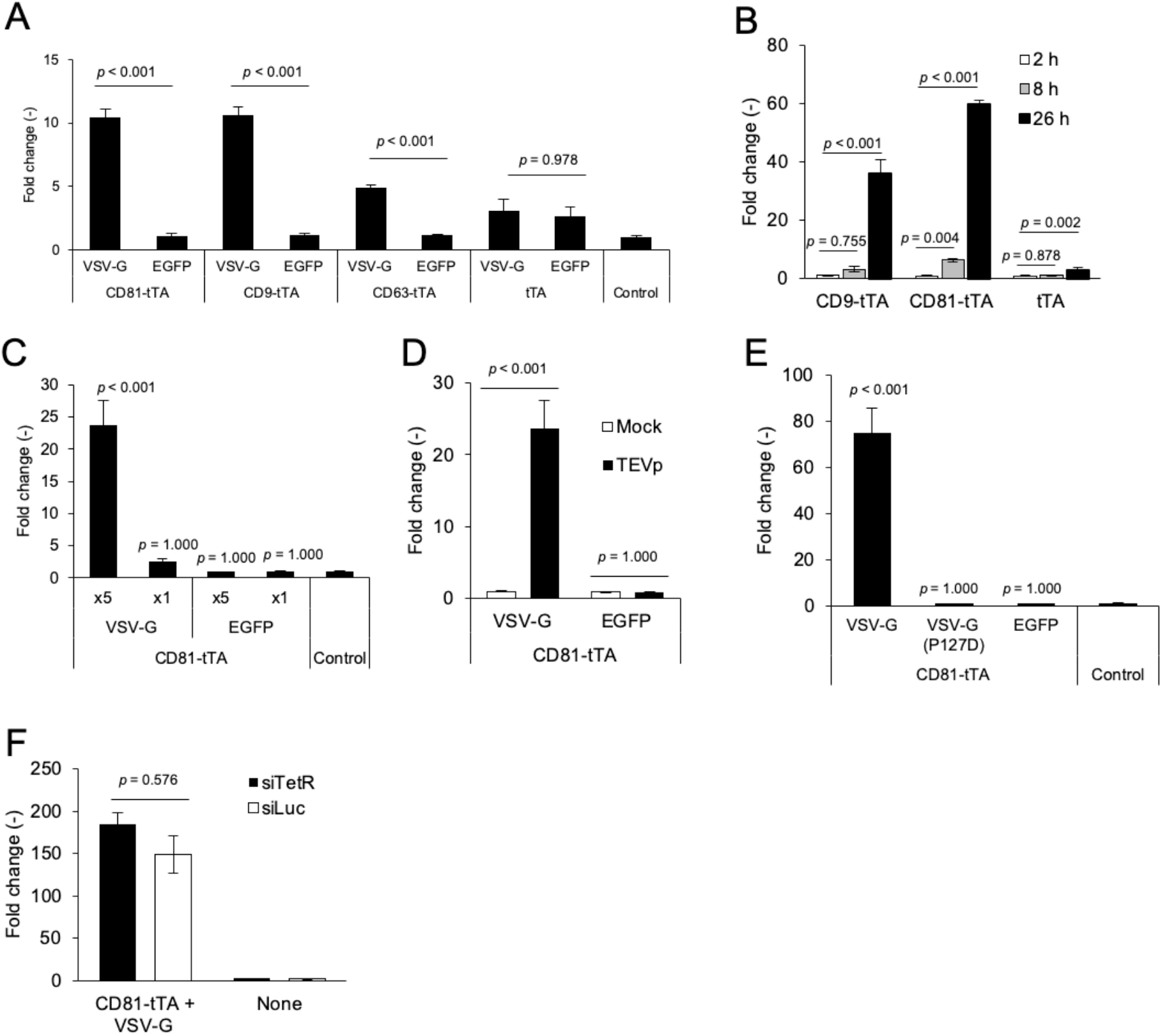
ETTD assay. (A) Donor supernatant was applied to recipient HEK293T cells and NanoLuc expression was measured after 24 h. NanoLuc expression was normalized to the control (treatment with supernatant from non-transfected donor cells). (B) Incubation time-dependent expression of reporter gene. Recipient cells were treated with concentrated VSV-G-expressing EVs containing tTA-fused CD9 or CD81 for 2, 8, and 26 h, followed by luciferase assay. (C) Dose-dependent reporter expression in recipient cells. Recipient cells were treated with EVs containing tTA-fused CD81 with or without VSV-G for 24 h. The relative amount of EV fraction added was noted as ×1 or ×5. (D) TEVp-dependent reporter gene expression. Recipient cells with or without expression of TEVp were treated with EVs containing tTA-fused CD81 with or without VSV-G and subjected to the luciferase assay after 24 h. (E) Effect of fusogenicity deficit VSV-G mutant. Recipient cells were treated with EVs (tTA-fused CD81) displaying parental VSV-G, mutant VSV-G (P127D), or EGFP for 24 h. (F) Recipient cells were pre-transfected with siRNAs targeting TetR (siTetR) or firefly luciferase (siLuc), and further treated with EVs containing tTA-fused CD81 and VSV-G for 24 h. N = 3, mean ± SD. Statistical analysis was performed using one-way ANOVA followed by *post hoc* Tukey’s HSD (A, B, D, E, and F) or Dunnett’s tests (C).

Next, we used EVs concentrated by PEG precipitation for the ETTD assay. Recipient cells were treated with EVs for 2, 8, and 26 h and the reporter NanoLuc expression was measured (Fig. 4B). NanoLuc expression gradually increased over time and reached a highest level at 26 h. The expression of NanoLuc was detected as early as 8 h. Induction of NanoLuc expression was dependent on the presence of VSV-G and the dose of EVs (Fig. 4C). Fig. 4D indicates that expression of TEVp in the recipient cells was crucial for reporter gene expression, demonstrating that the ETTD assay worked as expected (Fig. 1C). Furthermore, the EVs harboring fusion-deficient mutants of VSV-G (P127D)^19,22^ lost the membrane fusion ability of EVs in the assay compared to the EVs harboring parent VSV-G (Fig. 4E), strongly supporting that this assay depicted the membrane fusion-mediated cargo delivery of EVs. Furthermore, absence of VSV-G led to no functional delivery (Fig. 4C to 4E), indicating the poor cargo delivery efficacy of authentic EVs. In addition to HEK293T cells, HeLa cells were used as alternative recipient cells, and similar results were observed, indicating that the ETTD assay is potentially applicable to other cell lines (Fig. S1).

It was postulated that the excess of expression plasmid remaining in the supernatant or mRNA of tTA-fused tetraspanin encapsulated in EVs may induce the reporter gene expression in the recipient cells, which could confound the bona fide reporter expression due to the tTA release of EVs. Therefore, we transfected the reporter cells with siRNA targeting TetR, the TRE-binding domain of tTA to verify that the reporter gene expression was induced by tTA protein. First, we verified that siRNA targeting TetR (siTetR) efficiently knocked down tTA (Fig. S2A). Furthermore, knockdown of tTA by siRNA significantly suppressed TRE-mediated reporter gene expression (Fig. S2B). Based on these results, siRNA targeting tTA should abrogate the confounding factors in the ETTD assay, namely, the excess of expression plasmid remaining in the donor supernatant and mRNA-mediated expression of tTA. After transfection of siRNA into recipient cells, we applied tTA-fused EVs to recipient cells. As shown in Fig. 4F, transfection of siRNA targeting tTA showed no effect on the reporter gene expression, strongly suggesting that the assay readout of the ETTD assay was solely driven by tTA proteins, neither mRNA nor leftover plasmid DNA.

### Effect of small molecules on the membrane fusion efficiency of EVs

As the novel ETTD assay can evaluate the membrane fusion efficiency of EVs, we next examined the effect of potential delivery enhancers and entry inhibitors. According to a previous report, chloroquine enhanced Cre protein delivery of EVs by disrupting endosomes and lysosomes using the Cre-LoxP reporter assay ^14^. In our reporter assay, chloroquine treatment did not induce any reporter gene expression (Fig. 5A), suggesting that chloroquine does not enhance cytoplasmic cargo delivery of EVs. This is probably because chloroquine treatment induces the destabilization of endosomes/lysosomes and does not enhance membrane fusion.

**Fig. 5.**
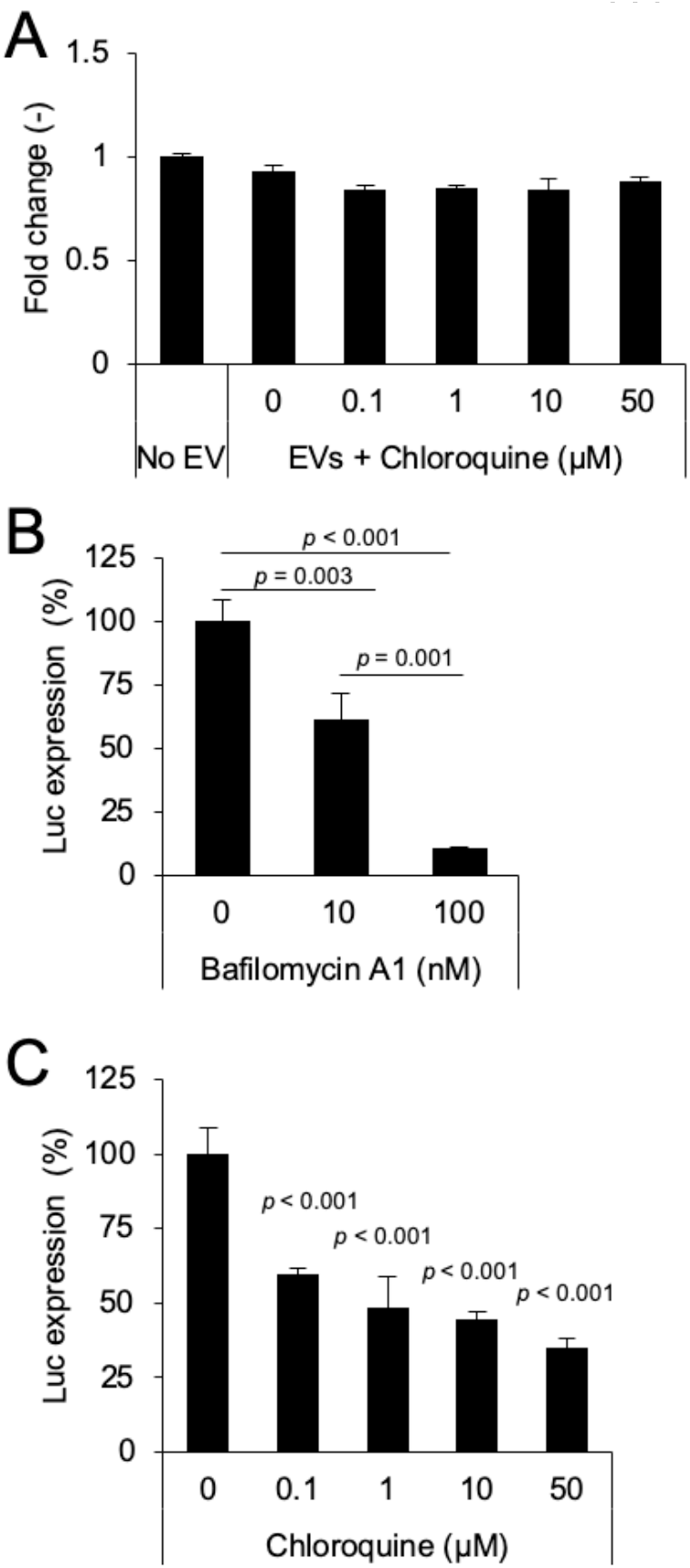
Effect of small molecule compounds on the ETTD assay. (A) EVs (CD81-tTA) without VSV-G were applied to recipient HEK293T cells in the presence of indicated concentrations of chloroquine. NanoLuc expression level was normalized to the value of the control (no EV treatment) and is presented as fold-change. (B) EVs (CD81-tTA) with VSV-G were applied to recipient cells in the presence of 10 or 100 nM of bafilomycin A1. (C) EVs (CD81-tTA) with VSV-G were applied to recipient cells in the presence of 0.1 to 50 μM of chloroquine. N = 3, mean ± SD. Statistical analysis was performed using one-way ANOVA followed by *post hoc* Tukey’s HSD test.

In addition to potential delivery enhancers, we assessed the effect of entry inhibitors using the ETTD assay. We used VSV-G-modified EVs to assess the effect of compounds that are known to increase the endosomal pH and thereby inhibit the low pH-dependent fusion activity of VSV-G ^23,24^. Bafilomycin A1 is a selective ATPase inhibitor ^25^ that prevents the acidification of endosomes/lysosomes and inhibits VSV infection ^26^. When recipient cells were treated with bafilomycin A1, membrane fusion by VSV-G-modified EVs was significantly inhibited in a dose-dependent manner (Fig. 5B). In addition, chloroquine, which is known to prevent VSV infection by increasing endosomal/lysosomal pH ^27^, also blocked the membrane fusion of EVs (Fig. 5C). These results strongly support the application of ETTD assay in assessing pharmacological effects of a potential delivery enhancer/inhibitor of EVs.

### Assessment of membrane fusion efficiency of EVs at the single-cell level

For the evaluation of EV membrane fusion at the single-cell level, we changed the reporter gene from NanoLuc to EGFP. As shown in Fig. 6A, EVs containing tTA-fused CD81 and VSV-G induced EGFP expression in the recipient cells, which was consistent with previous results (Fig. 4). This assay enabled us to decipher membrane fusion efficiency at the single-cell level.

**Fig. 6.**
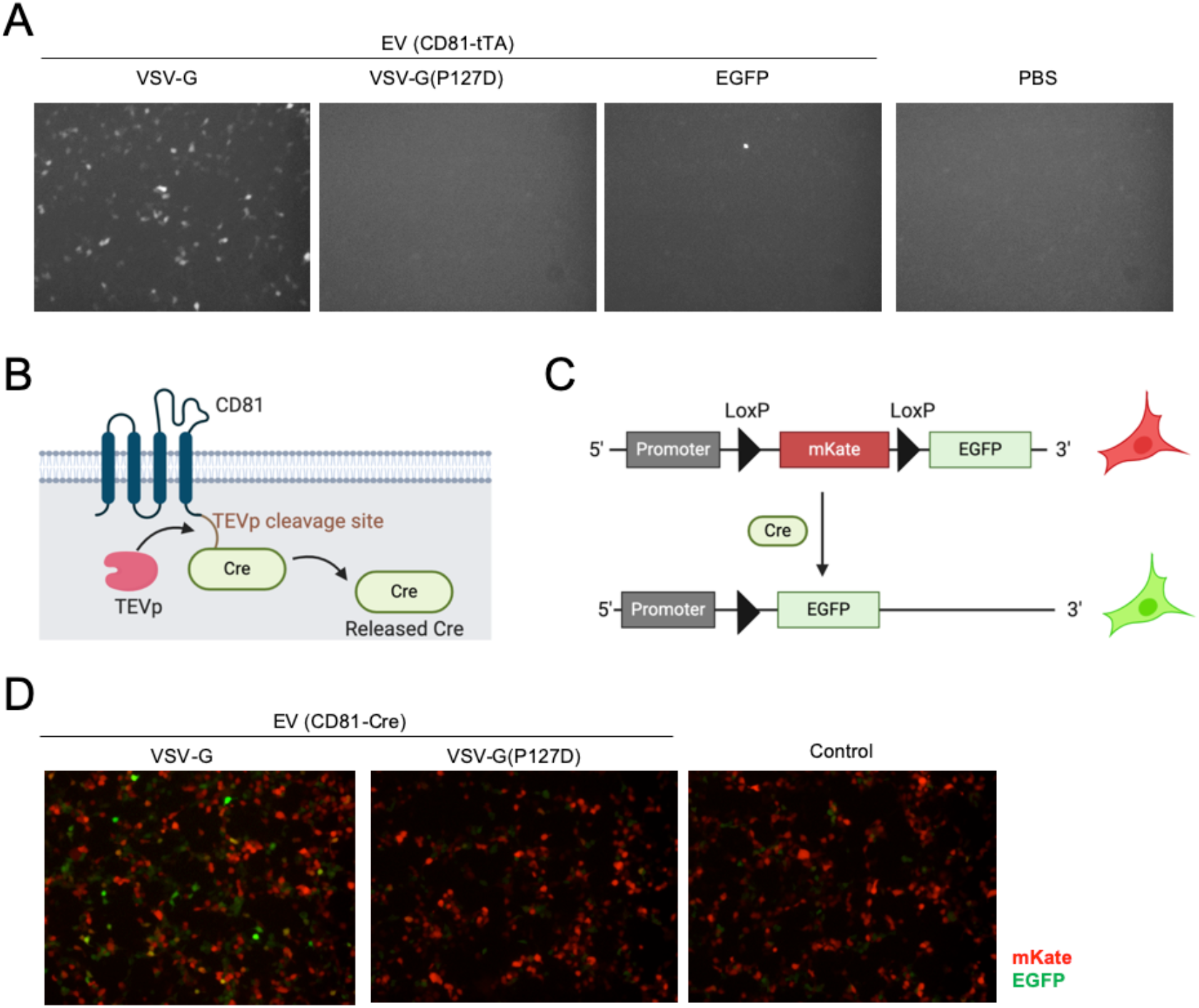
Fluorescence imaging-based reporter assays (A) EDDT assay based on EGFP as a reporter readout. Recipient HEK293T cells were transfected with plasmids encoding TRE3G-EGFP and TEVp, and treated with EVs containing CD81-tTA with VSV-G (WT or P127D mutant) or EGFP (negative control). After 24 h, cells were observed under a fluorescence microscope. (B) Schematic representation of Cre-fused CD81. (C) Schematic representation of reporter plasmid for Cre-LoxP reporter assay. After the Cre cleavage, mKate gene is excised and reporter cells become EGFP positive. (D) Fluorescence imaging of Cre-LoxP reporter assay. Recipient cells were treated with EVs harboring CD81-Cre and VSV-G.

To further validate the general applicability of the ETTD assay, we switched the reporter gene from tTA-dependent gene expression to the expression of a floxed gene dependent on Cre recombinase. The principle of the Cre-mediated reporter assay is essentially the same as that of the ETTD assay; however, the readout is driven by Cre-mediated recombination of the target gene. After the release of Cre from EV by TEVp, Cre recombinases translocate to the nucleus and induce recombination of the target plasmid (Fig. 6B & 6C). In this study, we used the mKate/EGFP reporter plasmid. The recipient cells initially expressed the red fluorescence protein mKate, but after Cre-mediated recombination, cells become EGFP positive (Fig. 6C). This assay may be more sensitive than the tTA reporter assay as even a single molecule of Cre recombinase can induce a readout in the recipient cells. As shown in Fig. 6D, EVs containing CD81-Cre with VSV-G induced EGFP positive cells, whereas EVs with VSV-G (P127D) showed almost no EGFP positive cells. This result was consistent with the results of the previous ETTD assay (Fig. 4) and again revealed that fusogenic proteins significantly increase the membrane fusion activity of EVs.

## Discussion

In this study, we developed an ETTD assay that can evaluate the membrane fusion efficiency of EVs in recipient cells. The principle of this assay was inspired by the tango assay ^28^ that quantitatively assesses receptor activation by the recruitment of genetically engineered TEVp to the receptor, subsequent release of tTA, and expression of TRE-mediated reporter gene. In the ETTD assay, tetraspanins constrain tTA and are localized at the membrane (Fig. 2B). Once the luminal tTA is exposed to the cytoplasm following membrane fusion of the EVs and release by TEVp cleavage, recipient cells express the reporter gene (Fig. 4D). This experimental design has rendered the ETTD assay robust and sensitive by avoiding non-specific background signals.

The ETTD assay enables us to quantitatively assess membrane fusion efficiency and delivery mechanism of EVs. The advantages of this assay are as follows: (1) it is highly sensitive to measure the membrane fusion of EVs with a wide dynamic range owing to the very bright NanoLuc, (2) fewer confounding factors in the ETTD assay compared to conventional assays because expression of the reporter gene under the TRE promoter is highly regulated and specific to the transcription factor tTA, which does not exist in mammalian cells; and (3) it is feasible to assess the membrane fusion efficiency in both the bulk cell population (NanoLuc reporter) and single-cell level (fluorescence protein reporter).

The very bright NanoLuc reporter gene, enables the ETTD assay to detect rare membrane fusion events in recipient cells. Because the cargo delivery efficiency of EVs is expected to be low (possibly 0.2% to 10% of recipient cells express reporter gene as a result of the cargo delivery, depending on the reporter assay ^15,29^), the assay sensitivity must be high to capture the membrane fusion events in recipient cells. When the EVs harbor the fusogenic protein, VSV-G, EV-mediated membrane fusion was sufficient for detection in the ETTD reporter assay, whereas no detectable membrane fusion was observed in the absence of VSV-G (Fig. 4). This result reflected the low efficiency of membrane fusion in the absence of a particular membrane fusion protein. As described in previous studies, the cargo delivery efficiency of EVs is expected to be low ^11,15,21,30–32^. However, our experiments were conducted using a combination of HEK293T donor cells and HEK293T or HeLa recipient cells. Other combinations of EV-donor cells and recipient cells should be examined to determine whether the cargo delivery efficiency is much higher in a future study.

We validated whether the ETTD assay precisely reflects tTA protein-mediated readout rather than mRNA transfer-dependent reporter gene expression. EVs can encapsulate overexpressed mRNA in the donor cells in a passive manner and potentially transfer the mRNA to recipient cells ^33^. Since it was postulated that unexpected EV-mediated transfer of tTA mRNA may lead to a false positive signal in the ETTD assay, recipient cells were pre-transfected with potent TetR-targeting siRNA (Fig. S2) and blocked the mRNA-mediated readout. The results clearly demonstrated that siRNA targeting TetR did not affect the assay readout, indicating the absence of mRNA-dependent tTA expression and subsequent reporter gene expression in the recipient cells (Fig. 4F).

Previously, membrane fusion of EVs has been evaluated by fluorescence probes ^34^ or reporter proteins ^19,20^. The former approach, especially the membrane-anchored fluorescence probes, such as R18, are known to often result in false positives due to non-specific dye transfer between lipid membranes ^35^. Joshi et al. developed a sophisticated fluorescence imaging technique to measure membrane fusion and cargo release of EVs in recipient cells ^36^. Their approach enabled the assessment of membrane fusion at the single-vesicle level; however, it was still difficult to distinguish the membrane fusion signal from the high background signal of the fluobodies distributed throughout the cytoplasm, and there was a limited capability in terms of throughput. The latter approach, typically using β-lactamase (BlaM) protein, is a time-consuming assay that requires a long incubation time for the enzymatic conversion of a fluorescence substrate (7 to 16 h ^19,20,37^). The ETTD assay, in contrast, is more feasible, sensitive, and rapid to assess the membrane fusion process of EVs in recipient cells and capable the high-throughput applications.

There are conflicting reports on the effect of chloroquine on EV cargo delivery in a previous study ^14^. In this study, chloroquine was unable to enhance membrane fusion and cargo delivery of EVs (Fig. 5A), whereas a previous study showed significant improvement in the Cre delivery of EVs. The inconsistency is probably due to the differences in the experimental settings, sensitivity, and accuracy between assays. The Cre-LoxP reporter assay is a sensitive and robust method since the assay readout is driven by ideally a single Cre molecule in the recipient cell, and assay readout is exclusively dependent on the Cre-LoxP excision of target DNA. The different conclusions between these studies should be carefully interpreted and further examined in a future study. Heath et al. demonstrated that small amounts of Cre recombinase (8.9 Cre-FRB molecules per EV on average) can be passively loaded into EVs and contribute to the recombination in the recipient cells ^14^, whereas our approach involved fusion of the Cre protein to the tetraspanin CD81 and application to the reporter recipient cells (Fig. 6B). It appears that our approach may be more convincing because the direct fusion of Cre with the EV marker protein is more reliable and precisely reflects the nature of EV-mediated cargo transfer.

## Conclusions

ETTD assay is a novel functional assay to assess the mechanism of EV-mediated membrane fusion and cargo delivery in a quantitative manner. The lack of reliable functional assays in the EV field has hampered progress in its therapeutic applications ^38^ and elucidation of the underlying mechanism of cargo delivery and intercellular communication of EVs ^10^. The ETTD assay is potentially useful for identifying unknown factors that are responsible for the cargo delivery mechanism. Using the ETTD assay, knockout or knockdown of target genes may reveal the unknown cargo delivery pathway as described in a previous study ^15^, or possibly facilitate the discovery of a methodology that enhances membrane fusion and subsequent cargo delivery of EVs. Together with the previously reported real-time cargo delivery assay ^21^, the ETTD assay may help advance fundamental EV research and its clinical applications.

## Acknowledgments

We extend our gratitude to Yumi Yukawa for technical assistance in plasmid construction. All illustrations in this work were created using BioRender.com.

This work was supported in part by JSPS KAKENHI (Grant-in-Aid for Early-Career Scientists 18K18386 and 20K15790 to MS), Research Grant from JGC-Scholarship (to MS), and the “Dynamic Alliance for Open Innovation Bridging Human, Environment and Materials” (MEXT).

