## Supplementary Data for "Reporter gene assay for membrane fusion of extracellular vesicles"

**Supplementary Methods**

*ETTD assay in HeLa cells*

Human cervical cancer-derived HeLa cells (JCRB Cell Bank) were used as recipient cells. Briefly, $10^4$ cells were plated in a 96-well plate, cultured overnight, and transfected with PEI and plasmids encoding TEVp and TRE3G-NlucP. The next day, the cells were treated with PEG-precipitated EVs and cultured for 24 h. Reporter NanoLuc expression level was measured as described in the main text.

*Co-transfection with siRNA and plasmid DNA*

HEK293T cells were co-transfected with siRNA and plasmid DNA using PEI. The antisense sequences of siRNA against firefly luciferase (siLuc) and TetR (siTetR) were 5'-UCGAAGUACUCAGCGUAAGtt - 3' [1] and 5'- UGAUCUUCCAAUACGCAACtt - 3' [2], respectively (lowercase letters indicate DNA). Briefly, $10^4$ cells were plated in a 96-well plate, cultured overnight, and transfected with PEI. PEI was mixed with 100 ng of plasmid and 1 pmol of siRNA. The weight ratio of PEI to pDNA was 4:1. The final concentration of siRNA used was 10 nM. Cells were cultured for 48 h and lysed to measure expression levels. For the quantification of HiBiT-tagged tTA, transfected cells were lysed and mixed with Nano-Glo HiBiT Lytic Detection System (Promega).

**Supplementary Figures**

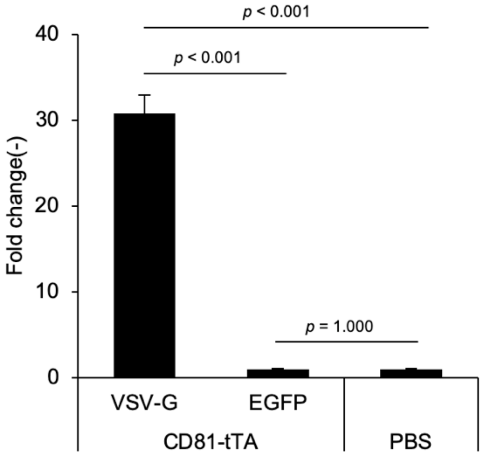


**Fig. S1 ETTD assay in HeLa cells**

Recipient HeLa cells transfected with TRE3G-NlucP and TEVp were treated with EVs containing tTA-fused CD81 and VSV-G or EGFP. PBS was used as a control.

N = 3, mean ± SD, Statistical analysis was performed using one-way ANOVA and *post hoc* Tukey's HSD test.

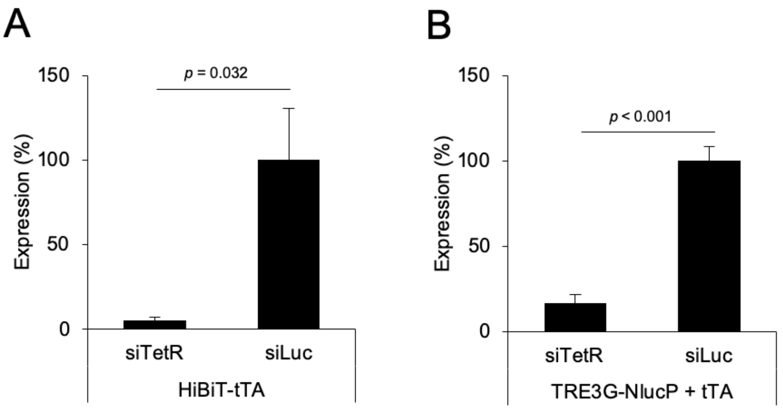


**Fig. S2 Knockdown of tTA by siRNA**

(A) HEK293T cells were co-transfected with HiBiT-tagged tTA and siTetR (targeting tTA) or siLuc (negative control) and expression level of tTA was evaluated by measuring HiBiT tag. The luminescence signal observed in siLuc sample was set as 100%.

(B) HEK293T cells were transfected with plasmids encoding TRE3G-NlucP and tTA, and either siTetR or siLuc. Expression level of NanoLuc was measured after 48 h and the luminescence signal observed in siLuc sample was set as 100%.

N = 3, mean ± SD, Statistical analysis was performed using Student's *t*-test.
